# Cartilage canals in sharks and rays show that blood vessels can exist in mature cartilage without triggering endochondral bone formation

**DOI:** 10.64898/2026.04.21.720020

**Authors:** Benjamin Flaum, Ronald Seidel, Maximus Yeatman-Biggs, Theda I. Hinrichs, Jana Ciecierska-Holmes, Salman O. Matan, Emilio J. Gualda, Kady Lyons, Victoria Camilieri-Asch, Susan R. McGlashan, Laura Ekstrom, Lawrence Bonassar, Mélanie Debiais-Thibaud, Daniel Baum, Michael J. Blumer, Mason N. Dean

## Abstract

Although cartilage in tetrapod skeletons is typically said to lack blood vessels, this is only true for adult cartilage. In young bird and mammal cartilage, a dense network of vasculature-containing tunnels —cartilage canals— perforate the growing skeleton, helping nourish the cartilage and develop the ossification centers that will later form the skeleton’s epiphyseal bone. As the canals and their rich vascular network typically recede as animals age, the healthy cartilage of adult animals is typically known to be avascular. Here, however, we use a range of tissue characterization and visualization techniques —including light/electron microscopy and microCT— to show that the skeletons of rays and sharks (elasmobranch fishes) not only possess cartilage canals, but that these structures persist in the adult skeleton. The morphology and tissue composition of elasmobranch cartilage canals argues homology with mammalian cartilage canals and an ancient invasion of the vascular system into cartilage. However, the anatomical location of canals —extending away from mineralized tissue not toward it— and the lack of endochondral ossification in ray and shark cartilage suggest that cartilage canals developed early in vertebrates as a transport system for nutrients and mesenchymal cells into the growing skeleton. We describe distinctive features and variation in elasmobranch cartilage canals, discuss their possible roles and their potential for tissue mineralization, and the biomedical implications for their presence in a clade of animals with continuously growing cartilaginous skeletons.

## Introduction

Most bones in the skeleton of tetrapod animals form indirectly from cartilage precursors converted into bone during early development and postnatal growth ^1,2^. During this process, known as endochondral ossification, the cartilage cells (chondrocytes) enlarge, triggering mineralization of the surrounding cartilage matrix as well as the invasion of blood vessels and bone-related cells into the cartilage scaffold. The cartilage matrix is then degraded by matrix metalloproteinases (MMPs), resorbed by chondroclasts and replaced by bone matrix (formed by osteoblasts), while the hypertrophic chondrocytes undergo cell death (apoptosis) ^3–8^. In the development of long bones (e.g. femur, tibia), a primary ossification center is first formed in the mid-shaft (diaphysis), followed by the formation of secondary ossification centers at the ends of the bone (epiphyses). Vasculature is key in this tissue transformation. Yet, whereas in developing primary ossification centers blood vessels invade the cartilage directly, in long bone secondary ossification centers (but also foot bones and vertebrae), blood vessels are ferried into the reorganizing unmineralized matrix via tunnels called cartilage canals ^9–11^. Cartilage canals are invaginations of the fibrous layer (perichondrium) that covers cartilage, migrating into the epiphysis via the action of matrix metalloproteinases, and vascular endothelial growth factor (VEGF, which attracts blood vessels) ^12–14^. In this way, cartilage canals allow a pocket of the skeleton’s external environment to penetrate the cartilage and contribute crucially to the formation of the localized ossification centers that establish the adult skeleton’s bone. However, the canals in tetrapods recede during development, replaced by the expanding bone tissue. As a result, cartilage canals and the vasculature they carry are considered absent in the cartilage of adult vertebrate animals.

Cartilage canals have been described in a range of terrestrial vertebrate species, especially mammals and reptiles (including birds) (see citations in Table S4). We show here, however, that cartilage canals —homologous to those of mammals— are also a primary component of the skeletons of adult rays and sharks (elasmobranch fishes), animals with skeletons made entirely of cartilage. Our results indicate that cartilage canals evolved far earlier than previously appreciated and, since the cartilaginous skeletons of these fishes are never converted to bone, that their cartilage canals have roles other than matrix ossification. Together, these inferences question the original role of cartilage canals in the vertebrate skeleton.

## Results & Discussion

### 1. In adult stingrays, as in young mammals, cartilage canals perforate the skeleton

The cartilage of rays, sharks and chimaeras is structurally unique among vertebrates in having a surface veneer of mineralized blocks called tesserae (Fig. 1C-F) ^15–17^. In rays and sharks, tesserae are typically hundreds of microns wide and deep, sandwiched between the outer layer of fibrous perichondrium and the unmineralized cartilage in the core of the skeleton (the latter grossly similar to mammalian hyaline cartilage) ^18–21^. The unmineralized cartilage, however, typically dominates ray and shark skeletons, with tesserae occupying a comparatively small proportion of skeletal cross-sectional area in most species (e.g. ∼10-20% in the Round stingray *Urobatis halleri*; ^22,23^).

**Figure 1.**
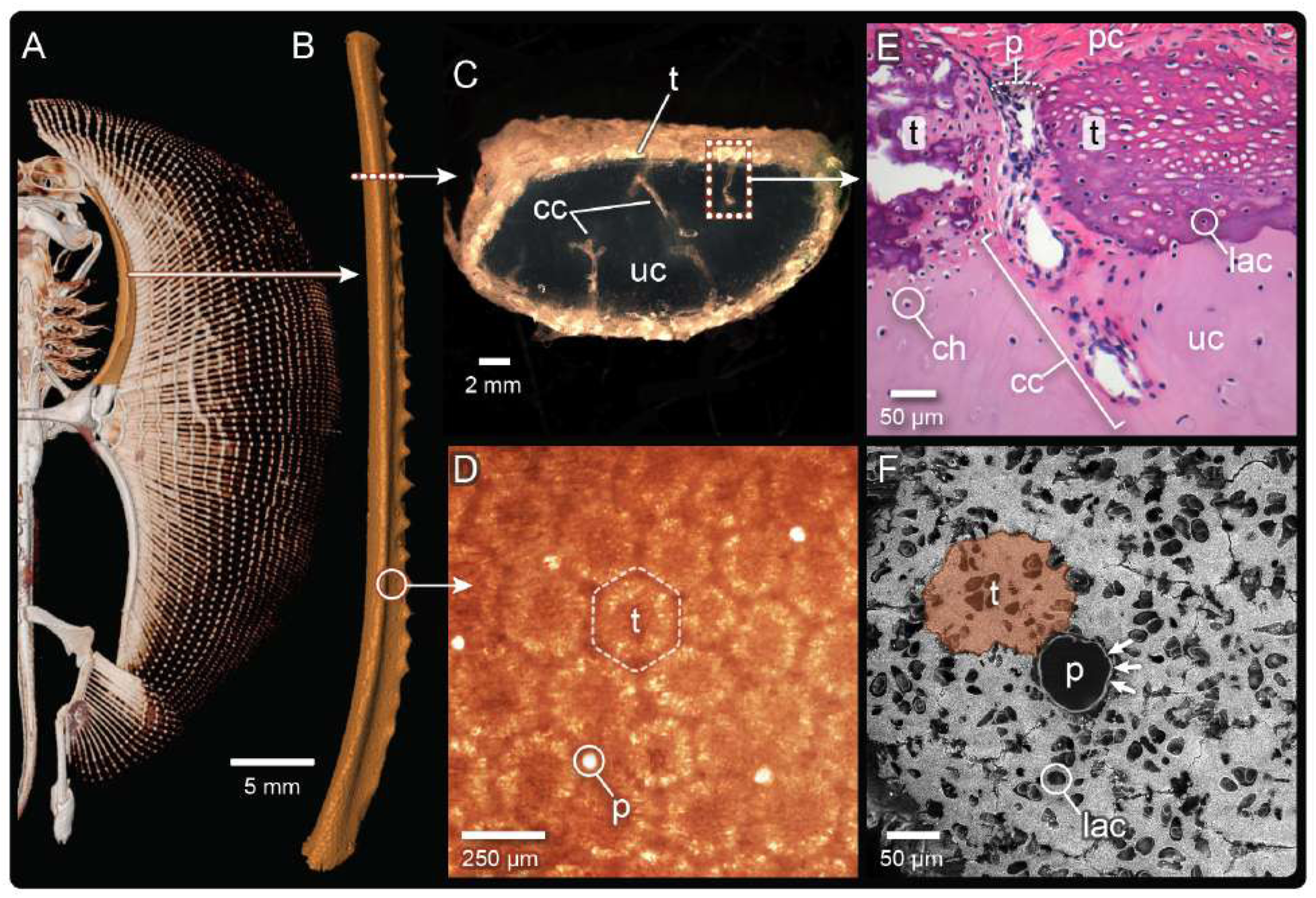
The Round stingray (*Urobatis halleri*, A), like all rays and sharks, has a skeleton made predominantly of unmineralized cartilage (uc) with surface mineralization. Individual skeletal elements (e.g. the propterygium, B), when cross sectioned (C), show an outer veneer of mineralized blocks (tesserae, t), which are polygonal when viewed from the skeletal surface (D,F). At the junctions of some tesserae, large pores (p) lead into cartilage canals (cc), which pass from the perichondrium (pc), between tesserae and into the underlying cartilage (D-F). The fibrous lining of canals/pores is visible in E (darker pink staining) and F (arrows), as are the chondrocytes (ch) within the unmineralized cartilage (E) and the lacunae (lac) within tesserae (F). Note that the darker pink staining around the canal in E is continuous with the fibrous perichondrium (see also Fig. 3C). A,B: µCT; C,D: light microscopy; E: H&E histology; F: BSE-SEM. All images from *U. halleri*, except C from *Leucoraja erinacea*.

We first observed cartilage canals in dissections of adult Round stingray(Fig. 1) ^24^, a small species we have used as a model for characterization of tesserae microanatomy and materials ^e.g. 18–21,25^. In this species, cartilage canals were visible with the naked eye in cross-sectioned specimens as white streaks or tubular gaps in the otherwise glassy cartilage matrix, but only once they had filled with a material of different refractive index than the surrounding matrix (e.g. blood or air, see Fig. S1). This optical behavior may explain why canals have almost entirely escaped notice in the anatomical literature (but see ^24,261–29)^. We characterized the cartilage canals of the Round stingray using a variety of tissue preparations and imaging approaches (e.g. unstained samples in light microscopy, stained histological samples, cryo-scanning electron microscopy); in diverse pieces of the skeleton; and in both sub-adult and adult specimens of varying ages (Table S3).

In the Round stingray, as in mammals, cartilage canals originate from the perichondrium and end blindly in the unmineralized cartilage core of the skeleton (Fig.1C,E); this basic topology was confirmed in many other elasmobranch species we examined (see below and Figs. 4, S3; Table S3). Passing from the perichondrium to the unmineralized cartilage, stingray canals did not bore through individual tesserae but rather traversed the mineralized layer via large pores (∼30-60 µm wide; Figs. 1D-F, S2, Table S1), found at the meeting points of multiple tesserae (Fig. 1D-F). Canal pores were irregularly spaced (515.7±144.3 µm apart in a 6.2 x 2.0 mm ROI, Table S1, Figs. 2A,S2) and strikingly obvious in light microscopy, electron microscopy (EM) and microCT preparations, being considerably larger and more circular than other joints or gaps between tesserae (Fig. 1D,F). Canal pores are therefore likely the intertesseral pores described by previous authors ^19,23,30–33^, although their role as conduits for cartilage canals was never realized.

**Figure 2.**
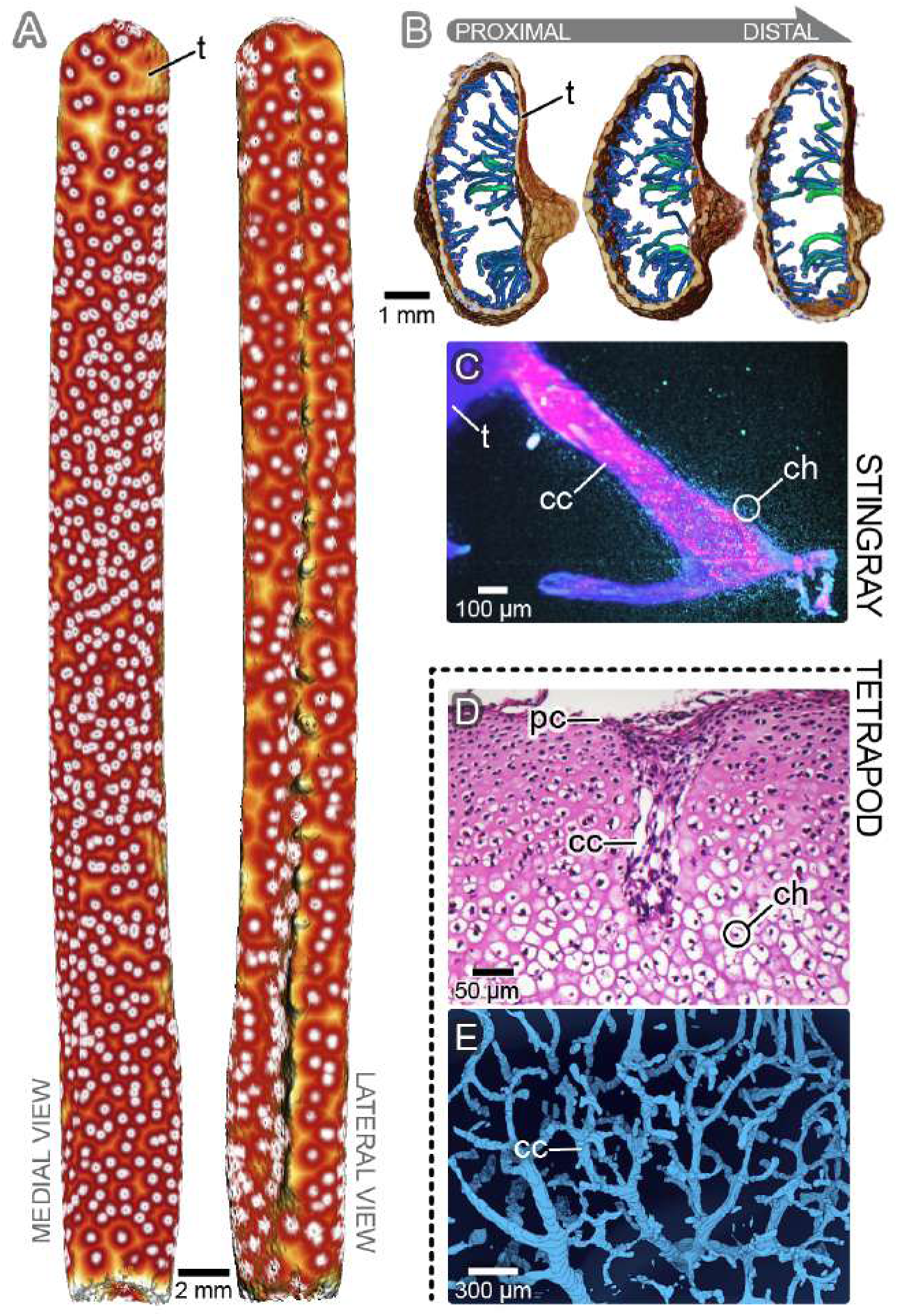
On the outer surface of the propterygium of the Round stingray (A), the origins of canals (marked by white dots) are densely arrayed on both the medial (left) and lateral (right) mineralized surfaces (tesserae, t). Skeletonized renderings of canals in three cross sections of the same propterygium (B), color-coded for diameter —using a gradient from blue (11 µm) to green (30 µm)— show canals that are largely similar in width but not length (see Table S2), with none passing entirely through the cartilage. Stingray cartilage canals (cc) are predominantly unbranched, but branching canals were also seen (C), and in all cases chondrocytes (ch) were found clustered tightly around canals in the matrix. In contrast, tetrapod canals (D,E) are only found in growing cartilage (D: note the high chondrocyte density in young D7 mouse epiphysis) and exhibit convoluted, dendritic networks (E: chicken e20 embryo epiphysis). A,B: quantitative µCT renderings; C: lightsheet microscopy (tissue autofluorescence); D: H&E histology; E: 3D reconstruction from histology (modified with permission from ^36^).

### 2. Stingray canals are prevalent, but simply structured and not linked to mineralization

Taking advantage of the fibrous lining of canals (see below), we treated a long piece of the wing skeleton with phosphotungstic acid (PTA), enhancing tissue contrast to make canals visible against the unstained cartilage in microCT scans (Fig. 2). We applied a custom semi-automatic segmentation approach, thresholding and digitally-isolating canals, allowing the entire population to be visualized and quantified (408 canals within a skeletal element ∼5 cm long and ∼4 mm wide/thick) (Fig. 2A,B). To our knowledge, no comparable quantitative 3D map of tetrapod canals exists, however images and 3D renderings from mammal and chicken cartilage show that canals are typically tens of microns in diameter and form a dense and richly dendritic network in the epiphysis, before the development of secondary ossification centers ^34–37^ (Fig. 2D,E). In tetrapod long bone epiphyseal cartilage, canals can be several millimeters long, extending sometimes nearly the full cartilage thickness to the deep hypertrophic zone of the epiphyseal growth plate ^34,36^. In contrast, stingray cartilage canals are far simpler in extent and form (Table S2), occupying only ∼0.3% of the cartilage volume, having a near-constant diameter (24.1±5.3 µm), similar to tetrapod canals, but being mostly unbranched (1.06±0.34 nodes), with bi- and trifurcated canals also rarely present (only ∼6% of canals) (Fig. 2B,C Table S2). Stingray canals were comparatively straight (1.08±0.08 tortuosity), running either parallel to the underside of tesserae or continuing perpendicular to them, extending deeper into the cartilage (Fig. 2B). In contrast to tetrapod epiphyses (e.g. Fig. 2D,E), no stingray canals traversed the entire cartilage bulk, instead ending <50% through the skeleton’s thickness (∼453.3±337.9 µm long; Fig. 2B, Table S2). Stingray canals are more densely distributed on average than those of newborn pup skeletons (571.42±176.79 µm vs. ∼1.4mm, respectively ^38,39^; Table S2), with apparent variability between lateral and medial surfaces of the propterygium, suggesting a degree of regionalization (Fig. 2A).

Despite the gross similarity between elasmobranch and tetrapod canals as ingress points into the cartilage bulk, their trajectories are notably different. In elasmobranch fishes, all margins of tesserae are thought to be active mineralization fronts, allowing tesserae to thicken and widen during development by accretion of new mineralized tissue layers ^15,18–20^. By perforating the tesserae layer to reach the uncalcified cartilage, elasmobranch canals therefore extend away from mineralizing zones rather than toward them, contrasting the association of tetrapod canals with ossification centers. This fundamental difference questions whether the canals of elasmobranchs and tetrapods are functionally and compositionally comparable.

### 3. Stingray canals are compositionally similar to those of tetrapods

As invaginations of the perichondrium, tetrapod cartilage canals are tubular conduits lined with perichondrial Type-1 collagen, penetrating deep into the mineralizing cartilage matrix, carrying mesenchymal cells, unmyelinated nerve fibers, macrophages, TRAP-positive cells (e.g. chondroclasts), and vasculature embedded in loose connective tissue ^14,34,39–41^. Similarly, our histology and EM preparations of stingray cartilage show a collar of fibrous material lining the canals (Fig. 3) and immunohistochemistry (IHC), verified that this is Type-1 collagen, whereas the surrounding cartilage matrix is patterned on a cartilage-typical Type-2 collagen (Fig. 3C-F). We saw no evidence of nerve fibers or TRAP-positive cells (e.g. breaking down the tesserae’s mineralized matrix). Importantly, however, stingray cartilage canals contain capillaries enmeshed in loose connective tissue (Figs. 3B,D,F), debunking the long-standing beliefs that the elasmobranch skeleton is avascular and/or inhibits vascular invasion ^1,16,42,43^. Adult stingray canals therefore, surprisingly, mirror canal composition in juvenile tetrapods, representing a perichondrial microcosm that provides entrance for vasculature into the skeleton. This demonstration challenges the textbook dogma that adult healthy cartilage lacks vasculature, suggesting an entirely different physiological environment for the adult cartilage of stingrays relative to that of other vertebrates.

**Figure 3.**
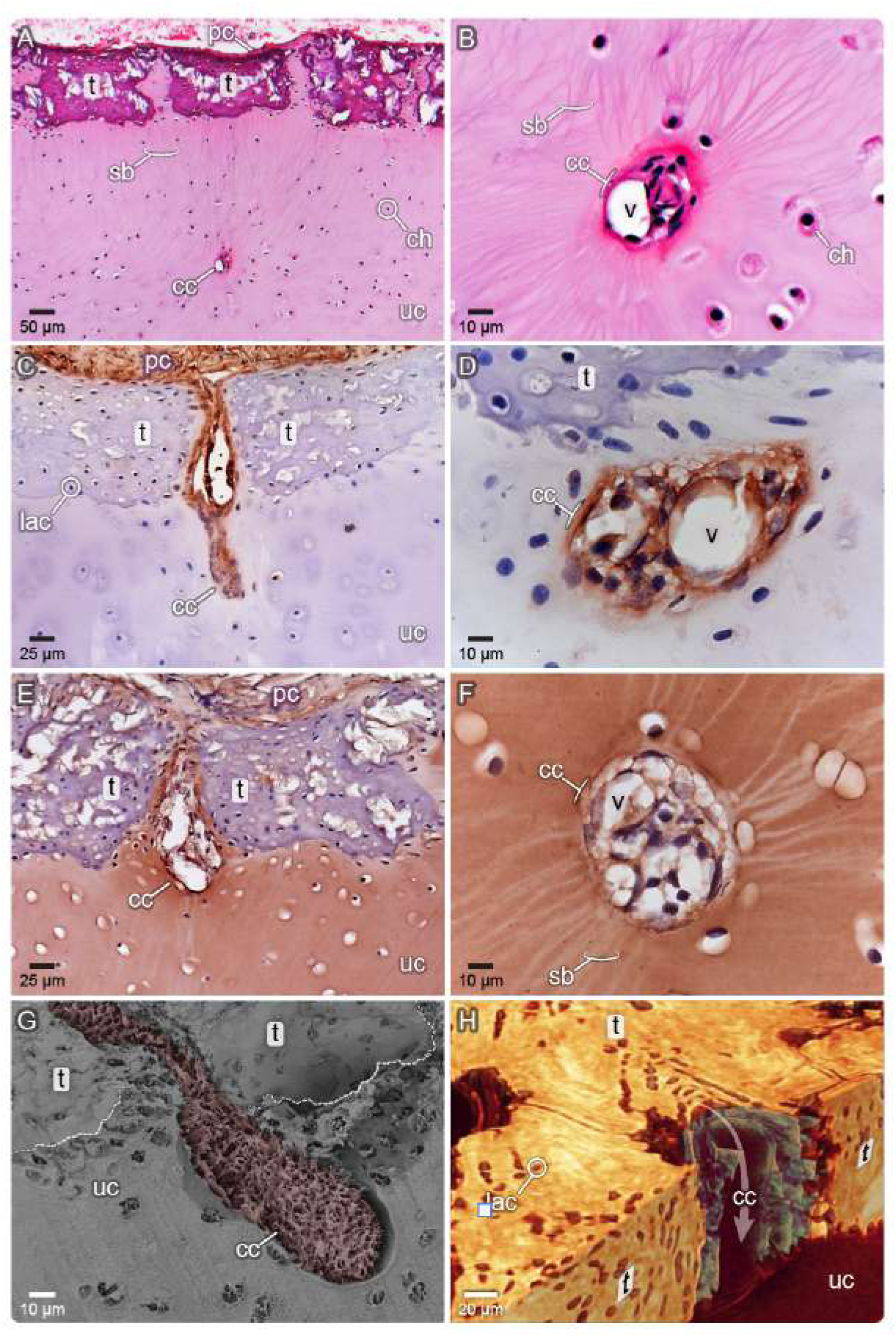
In cross section within the unmineralized cartilage (uc) matrix, cartilage canals (cc) are roughly circular (A,B,D,F), with chondrocytes (ch) and threadlike Saftbahnen (sb) visible surrounding them, and containing vessels (v). Immunohistochemistry (brown staining) shows that canals are invaginations of Type-1 collagen (stained brown in C,D) from the perichondrium (pc), whereas the unmineralized cartilage matrix is Type-2 collagen-based (E,F). The canal’s lining is therefore structurally distinct from the surrounding cartilage matrix, as shown in a freeze-fractured sample (G) of a longitudinally-sectioned canal with the fibrous lining intact (i.e. the Type-1 collagen layer is viewed from its peripheral aspect, the canal lumen and its contents are not visible). A virtual section of µCT data (H) —with tesserae (t) sectioned in-plane on the upper left and a canal and two tesserae sectioned vertically on the bottom right— illustrates how the orientations of cell lacunae (lac) track the shape of canals in two planes (see also ^30^). A,B: H&E histology; C,D: Type-1 collagen IHC; E,F: Type-2 collagen IHC; G: cryoSEM with canal lining pseudocolored; H: µCT with tesserae, canal lining, and other non-mineralized background (including lacunae within tesserae) pseudocolored yellow, green, and red, respectively.

### 4. Cartilage canals are widespread features in shark and ray skeletons

To verify that cartilage canals were not unique to the Round stingray among elasmobranch fishes, we assembled observations from a broad phylogenetic range of shark and ray species, many from specimens collected for other studies and which provided skeletal cross sections for evaluation (Table S3, Figs. 4, S3). As a result, age could not be controlled for and sampling location was not consistent, but tended to be primarily in the visceral arches (jaw, hyoid, branchial arches), but also the pectoral/pelvic girdles and chondrocranium. Given that multiple canals were typically visible in each skeletal cross section from the Round stingray, we coded species as having canals if any (i.e. ≥1) could be definitively observed across several specimen sections. Larger canals that pass through the vertebral body of some large sharks were excluded ^44^, as they run only through a thick, highly-mineralized tissue called ’areolar cartilage’ ^45–47^.

Whereas cartilage canals were identified in all 17 examined species of batoid fishes (rays, skates, guitarfishes), canal presence was inconsistent in the 17 shark species investigated, with some families having species with and without canals. Additionally, canals were not observed in chimaera, lamprey or hagfish, species included as outgroup comparisons. Allocation of certain species as lacking canals is not a definitive demonstration of canal absence in those elasmobranch families; further systematic evaluation is required. However, the variation in the frequency of canal occurrence suggests that canals may be more physiologically important in some groups. Concomitantly, canals also showed a range of morphologies: whereas most batoid species had canal morphologies similar to the Round stingray’s, two sharks from different families (Common thresher *Alopias vulpinus* and Sandbar shark *Carcharhinus plumbeus*; Fig. 4C, S1) had short and squat canals, while angel sharks (*Squatina* spp.) had particularly elaborate canal systems (Fig. 4B). The suggested phylogenetic signal within the variation in cartilage canal network architecture also points to a potential diversity in canal role and function.

**Figure 4.**
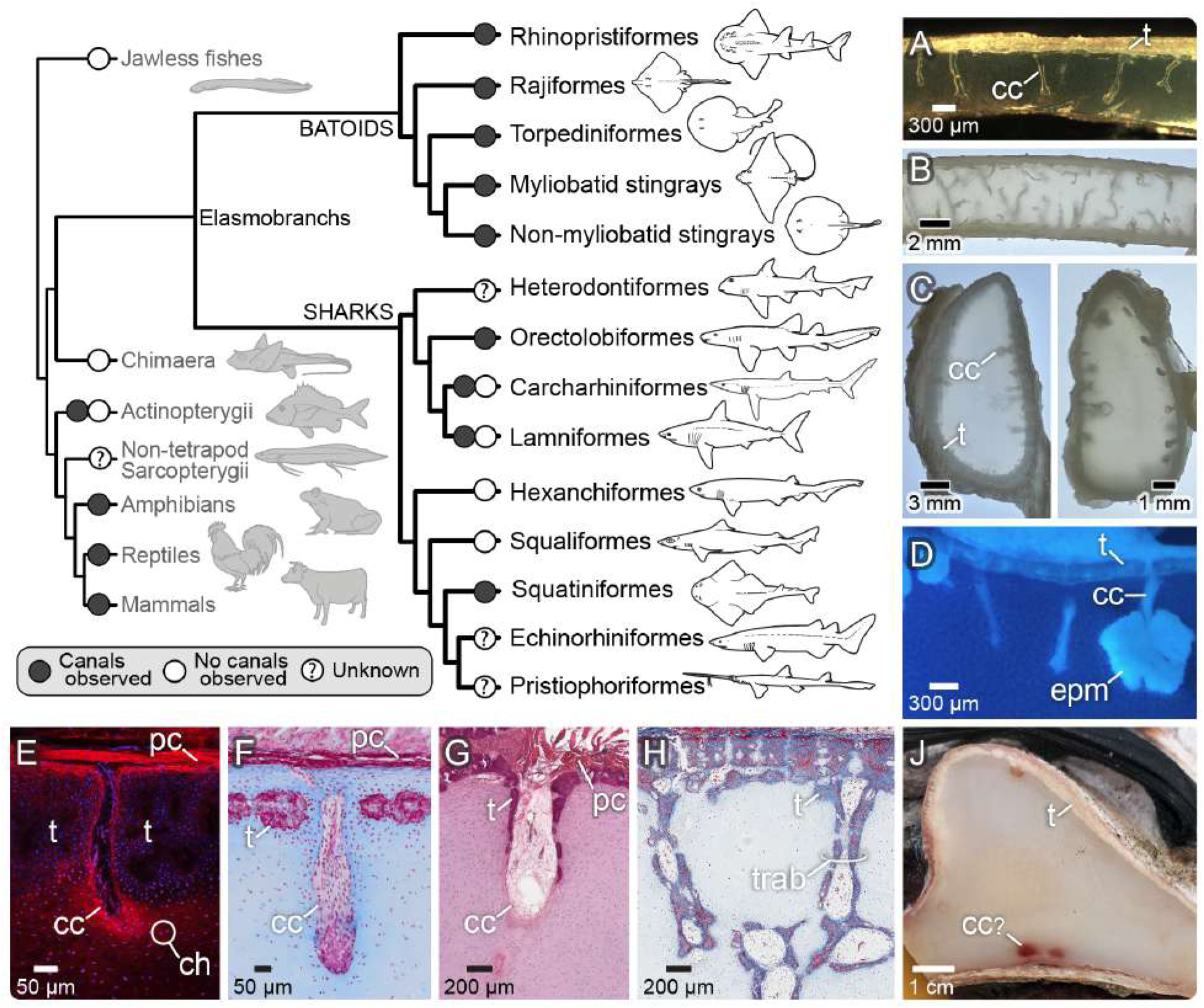
A phylogenetic survey of cartilage canal presence in elasmobranch fishes suggests that they are widespread in batoid fishes, but more patchily distributed in sharks. Combined with our data for outgroups (chimaera, jawless fishes) and literature reports for other vertebrates (see Tables S3,S4), current data suggest cartilage canals arose before or at least without a link to endochondral ossification. Circles at branch tips indicate whether canals were observed in the elasmobranch family members we examined or have been reported in the literature for any species of a group (see Tables S3,S4,); question marks indicate groups for which no data currently exists. Images of cartilage canals (cc) from various elasmobranch species (A-J, Table S3) illustrate that all instances show a similar arrangement, extending from the perichondrium (pc) between tesserae (t) to end in the unmineralized cartilage. Species, however, show a wide morphological variation, including sparsely branched canals similar to those in Round stingray (A: Batoid, *Leucoraja erinacea*); denser multibranched canals (B: Squalean shark, *Squatina squatina*); and short bulbous canals (C: Galean sharks, *Isurus oxyrinchus* (left) and *Carcharhinus plumbeus* (right)). Endophytic masses (epm) were seen associated with some canals (D: Batoid, *Urobatis halleri*) and chondrocytes (ch) clustered around all canals examined at histological resolutions (e.g. E: Batoid, *Mobula tarapacana*; F: Galean shark, *Galeorhinus galeus*). In some species, canals show varying degrees of lining by tesserae, from partial coverage (G: Batoid, *Dasyatis pastinaca*) to fully-covered trabeculae (trab) (H: Batoid, *Rhinoptera bonasus*). In the samples we had from some species, no canals were observed (see Table S3); in the case of the basking shark (J: Galean shark, *Cetorhinus maximus*), canal presence was ambiguous, despite the massive thickness of the cartilage. See Fig. S3 for additional images of canals from examined species not shown here.

### 5. Implications

Tetrapod cartilage canals have three major putative functions. First, canals nourish growing epiphyseal cartilage, which would otherwise depend exclusively on passive diffusion of nutrients from capillary networks external to the skeleton ^48^. Perfusion examinations ^38^ revealed that each canal contains a central arteriole that branches into a capillary glomerulus from which venules return to the perichondrium, thereby supplying the cartilage with nutrients and eliminating waste products. At the same time, an interruption of the cartilage canals’ blood supply by minor trauma can be an initiating factor of osteochondrosis, where ossification of the epiphysis is impaired and necrotic islands of cartilage are retained ^49–51^. Second, several studies suggest that canals contain Type-2 collagen-synthesizing cells, which are responsible for the interstitial growth of the epiphysis ^34,40,48,52,53^. In piglets and foals, this occurs via chondrification, where canals regress at their distal ends, their endothelial cells apoptose, and their fibroblasts transdifferentiate into matrix-producing chondrocytes ^49–51,54,55^. Lastly, fibroblasts and mesenchymal cells in the connective tissue filling canals also express several bone-related proteins (Type-1 collagen, periostin, alkaline phosphatase, and Runx2), suggesting that they may later differentiate into bone cells (osteoblasts and osteocytes), which are crucial for the mineralization of the organic bone matrix (Type-1 collagen) and thus formation of the secondary ossification center ^9,48,56–58^.

The vast majority of cartilage canal research has focused on only a few species of mammals and birds (Table S4), where the canals have very similar structural characteristics. Mapping the presence of canals following the limited published species data (Tables S3,S4) onto the vertebrate phylogeny suggests that canals either arose independently several times (in elasmobranch fishes, tetrapods and in sturgeon) or once at the base of jawed vertebrates (gnathostomes) and then were patchily lost within clades (Fig. 4). Within elasmobranchs, our demonstration of cartilage canals in a wide diversity of lineages suggests they are an ancestral feature for the clade with high potential for secondary loss (and maybe secondary gains). With our current knowledge, we consider the pervasive presence of cartilage canals in elasmobranchs and tetrapods, with shared histological features, to be the result of a single evolution in their last common ancestor. This means that one of the major anatomical features that permit conversion of cartilage into bone was already present in stem vertebrate lineages but was likely only later co-opted into an ossification function in tetrapods. Future thorough sampling within clades —especially the hugely speciose actinopterygian bony fishes—will be vital for evaluating the phylogenetic extent of cartilage canal presence, to help understand the role cartilage canals played in animal habitat radiation and skeletal tissue evolution.

Since chondrocytes rely on diffusion of nutrients through the gel-like extracellular matrix, it is thought that they must remain local to vasculature; in newborn canine pup skeletons, chondrocytes are never more than ∼700 µm away from canals ^38,39^. This constraint imposes profound limits on both the cell density and the size of cartilage elements in adult tetrapods, where healthy cartilage typically lacks blood vessels: in the femoral condyles of land tetrapods, articular cartilage thickness increases with negative allometry relative to body mass, with the largest species studied (e.g. elephant, rhinoceros, horse) having cartilage just ∼2-3 mm in thickness, with cell density decreasing greatly the further away from articular surface ^52,59,60^. There are no comparable datasets spanning the diversity of sharks and rays, but our dissections of various species regularly revealed skeletal cross sections centimeters in thickness, with some specimens of Basking shark (*Cetorhinus maximus*) cartilage exceeding 4-5 cm thick (oddly, however, canals could not be unequivocally demonstrated in this species; Fig. 4J). Although a cartilage skeleton is a unifying characteristic of all sharks and rays, elasmobranch fishes are diverse, with >1200 species exhibiting a rich range of ecologies and habitats and a huge range of body sizes, from tens of centimeters to >10 m in length ^61–63^. The presence of canals may have released a maximum size constraint ^63^ on elasmobranch cartilaginous skeletal elements: most canals we observed did not pass completely through the cartilage bulk, yet with the perichondrium surrounding the entire skeleton (unlike mammalian epiphyses cartilage), canals can enter —and supply tissue— from any direction.

Elasmobranch cartilage canals are tightly associated with chondrocytes. Where canals pass into the skeleton, cells within tesserae are oriented to follow the canal’s path (Fig. 3H) and are in significantly higher density than regions not bordering pores ^30^. Our lightsheet microscopy and histology (Fig. 2C, 3A) also show high densities of chondrocytes haloing canals within the unmineralized cartilage; as a result, the location of canals could be identified from adjacent histological serial sections, even when the canal itself had not been captured in the section. Previous work on the little skate (*Leucoraja erinacea*, a batoid species) demonstrated that progenitor cells originating in the inner layer of the perichondrium were able to traverse cartilage canals and differentiate into chondrocytes deep within the unmineralized cartilage ^27^. Additionally, we observed strand-like structures (1-2 µm diameter) radiating from canals (Fig. 3A,B,F) —visible with multiple staining and imaging techniques— often connecting to the basal surfaces of tesserae and linking chondrocytes hundreds of microns away in a daisy-chain series. These odd structures —variously-termed “*Saftbahnen*” or “interlacunar networks”— have also been observed in chimaera, chicken, mouse and rat cartilage ^see^ ^citations^ ^in^ ^21^, yet their function remains unknown and understudied. Their association with cartilage canals in elasmobranchs, however, indicates that *Saftbahnen* evolved early in vertebrates perhaps in concert with canals, and therefore may be linked to chondrocyte nourishment, cartilage growth and, later, mineralization.

Although matrix mineralization is clearly not a primary role for elasmobranch cartilage canals, our findings suggest they may still harbor the potential to promote calcification. In specimens of several different species, we observed endophytic masses within the cartilage (Fig. 4D); these disordered mineralized features are hypothesized to result from damage or aberrant mineralization processes in elasmobranch skeletons ^22^. Our observations of endophytic masses attached to some canals suggest the two features may develop dependently. Similarly, in several cases, we observed tesserae that had formed along the proximal ends of canals, tracking them for short distances along their dive into the matrix (Fig. 4G). Myliobatiform stingrays are an extreme case of this phenomenon where long trabeculae, lined entirely by tesserae, form reinforcing struts in the jaws, mostly in species that eat hard-shelled prey (Fig. 4H) ^23,64,65^. We have found the lumen of these trabeculae contains similar tissues to cartilage canals (Fig. 4H), indicating that canals may have been structural precursors to trabeculae, therefore arguing they also played a role in the evolution of specialized diets in elasmobranchs.

Our discovery of persistent vascular cartilage canals as widespread features in the cartilage of adult sharks and rays raises a host of further questions regarding their ubiquity in elasmobranchs and other taxa, development and evolution, and their physiological activity. Understanding the metabolic and mechanical factors that regulate the growth, invasion in cartilage, and persistence of these canals (i.e. the lack of endochondral ossification) should give clues to how some elasmobranchs attain massive skeletal masses and exceptional lifespans (likely the longest among vertebrates ^66^). In modern orthopedic medicine, the repair of joint cartilage poses a major challenge, primarily because it lacks a blood supply and therefore has only limited self-healing capacity ^67,68^. When devitalized mammalian articular cartilage plugs are grafted to vital cartilage plugs or implanted subcutaneously in nude mice, soft tissue canals can develop from the vital tissue, penetrating devitalized tissue ^67–70^. As cartilaginous fish skeletons continue growing throughout life, the study of their cartilage canals may provide keys to further biomedical methods for producing responsive implants for damaged cartilage, as well as immortalized (infinitely proliferative) cell lines and cutting-edge treatments for joint diseases.

## Acknowledgments

The authors thank Ocean Park Aquarium for donating specimens, Cheryl Wilga for generously sharing hard-won images of shark cartilage, and Daniel Fernando, Akshay Tanna, and team at Blue Resources Trust (BRT, Sri Lanka), who facilitated access to mobulid specimens from local fisheries and their export to QUT in Australia. Julien Lesseur, Daniel Werner and Júlia Chaumel helped arrange the fantastic microCT data for Fig. 2. Dominique Adriaens and Annabella Knab generously helped with histology. We thank the core facilities at Cornell University, specifically the College of Veterinary Medicines’ Animal Health Diagnostic Center (AHDC) for embedding and staining the *R. bonasus* sample and the Aperio ScanScope Unit for imaging it. Additionally, we thank CityUHK Veterinary Diagnostic Laboratory for embedding and staining the *A. ocellatus* sample. Peter Fratzl and Adam Summers, as always, provided support and sharpened curiosity that drove the project forward, as did the lovely community of comparative skeletal biology researchers at the IAFSB/CCB/CCBB conferences over the past decade, since this project began.

## Author contributions

Benjamin Flaum: Conceptualization, Methodology, Validation, Formal analysis, Investigation, Resources, Data Curation, Writing - Original Draft, Writing - Review & Editing, Visualization, Project administration, Funding acquisition

Ronald Seidel: Conceptualization, Methodology, Software, Validation, Formal analysis, Investigation, Data Curation, Writing - Review & Editing, Visualization

Theda Hinrichs: Methodology, Validation, Formal analysis, Investigation, Resources, Data Curation, Writing - Review & Editing, Visualization

Maximus Yeatman-Biggs: Methodology, Validation, Formal analysis, Investigation, Resources, Data Curation, Writing - Review & Editing, Visualization

Jana Ciecierska-Holmes: Validation, Formal analysis, Investigation, Writing - Review & Editing, Visualization

Salman O Matan: Validation, Investigation, Writing - Review & Editing

Emilio Gualda: Methodology, Validation, Formal analysis, Investigation, Resources, Writing - Review & Editing, Visualization

Kady Lyons: Investigation, Resources, Writing - Review & Editing

Victoria Camilieri-Asch: Methodology, Validation, Resources, Writing - Review & Editing, Project administration

Sue McGlashan: Methodology, Validation, Resources, Writing - Review & Editing, Supervision, Project administration, Funding acquisition

Laura Ekstrom: Methodology, Validation, Resources, Writing - Review & Editing, Visualization

Lawrence Bonassar: Methodology, Validation, Resources, Writing - Review & Editing, Supervision, Project administration, Funding acquisition

Mélanie Debiais-Thibaud: Conceptualization, Methodology, Validation, Formal analysis, Investigation, Resources, Data Curation, Writing - Original Draft, Writing - Review & Editing, Visualization, Supervision

Daniel Baum: Conceptualization, Methodology, Software, Validation, Formal analysis, Investigation, Resources, Data Curation, Writing - Review & Editing, Visualization, Supervision

Michael J. Blumer: Conceptualization, Methodology, Validation, Formal analysis, Investigation, Resources, Data Curation, Writing - Original Draft, Writing - Review & Editing, Visualization, Project administration

Mason N. Dean: Conceptualization, Methodology, Validation, Formal analysis, Investigation, Resources, Data Curation, Writing - Original Draft, Writing - Review & Editing, Visualization, Supervision, Project administration, Funding acquisition

## Star Methods

### 1. EXPERIMENTAL MODEL & STUDY PARTICIPANT DETAILS

The majority of data for this work is reanalyzed from previous studies (see Table S3 for sources); the following are the additional specimens examined specifically for this study:

#### Urobatis halleri specimens

Three Round stingray (*U. halleri*) specimens were used in this work, all donated from a previous study where they were collected in Seal Beach, California, USA ^71^. The tissues described below were excised and kept for the current study, while other portions of the skeleton were used for other previous studies (e.g. ^18,19,21,22,24,25^).

First, the left propterygium from a 21.1 cm disc width (DW) male was excised and used for phosphotungstic acid (PTA)-stained µCT scanning. Second, tissue from an 11.5 cm DW male was used to quantify pore morphometrics (Fig. S2). A flat region of tesserae (∼6.2x2.0 mm) was excised from the side of the propterygium and most of the unmineralized cartilage removed, leaving predominantly the tessellated layer. Lastly, a cross section of propterygium from an 11.0 cm DW male was used to examine how canal visibility changes with tissue dehydration (Fig. S1).

#### Devil ray (Mobula) specimens

Specimens of several devil ray species (*M. mobular*, *M. thurstoni*, and *M. tarapacana*) were obtained from Sri Lankan target fisheries under CITES permit 23SL003352, and university ethics approval for tissue use (QUT2000000191). The *M. eregoodoo* specimens were provided as bycatch from the Queensland Shark Control Program under general fishing permit 213613, standard goods BICON permit 0007233697, and DAFF Case ’Preserved and fixed animal and human specimens’ (effective: 5 April 2023)

#### Galean and squalean shark specimens

Copper shark (*Carcharhinus brachyurus*), school shark (*Galeorhinus galeus*), blue shark (*Prionace glauca*) and greeneye spurdog (*Squalus chloroculus*) specimens were obtained as commercial bycatch off Mangawhai (*C. brachyurus*, *G. galeus)*, East Cape (*P. glauca*) and Whangaroa (*S. chloroculus*), New Zealand.

#### Eagle ray (Aetobatus ocellatus) and cownose ray (Rhinoptera bonasus) specimens

The eagle ray (*A. ocellatus*) specimen was donated to the project from Ocean Park Hong Kong. The cownose ray (*R. bonasus)* specimen was bycatch from fishermen off of North Carolina, USA.

### 2. METHOD DETAILS

#### Transmitted light microscopy

To measure pore morphometrics, a transmitted light microscopy image of the isolated tessellated layer of a Round stingray specimen (11.5 cm DW) was reduced to 8-bit, thresholded (Otsu, 0-250), inverted, eroded and dilated using Fiji ^72^. Python code, using *skimage.morphology.remove_small_objects*, removed noise. *Skimage.measure.label* and *.measure.regionprops* were used to label and extract the maximum Feret’s diameters of 28 pores ^73^. Centroid coordinates and *scipy.spatial.KDTree* were used to locate the nearest neighbor, and *scipy.spatial.distance.cdist* to calculate minimum Euclidean distance between pore edges ^74^.

To examine how canal visibility changes with dehydration, the propterygium cross section from the 11.0 cm DW animal was placed in a drop of water in a Petri dish and observed under a Keyence Digital Microscope. Images of the entire cross section were taken sequentially as the sample dehydrated and a canal in the uncalcified matrix became apparent.

#### Phosphotungstic acid-stained microcomputed tomography (µCT)

The left propterygium of a Round stingray *U. halleri* specimen (21.1 cm DW) was excised and fixed in 75% ethanol. The propterygium was soaked in a 1% PTA solution for 36 hours. Post-staining, the whole skeletal element was scanned in an RX Solutions EasyTom160 microCT scanner (Chavanod, France) at 80 kV source voltage and 125 µa source current with a final voxel size of 11 µm.

Reconstructed image slices were imported into Amira ZIB software ^75^ where the data were downsampled to 22 µm voxel size and the cartilage canals were segmented (digitally dissected) semi-automatically using the magic wand tool, producing a label field of the canal network. An *Autoskeleton* module was attached to the resulting label field, from which all morphometric data were extracted. A Euclidean distance map was applied to the thinned label field from the *Autoskeleton* module, from which the distances from each canal to all others were determined, with the shortest pairwise distance for each canal taken as the nearest neighbor distance.

#### Unstained microcomputed tomography (µCT)

µCT data of air-dried, unstained Round stingray cartilage was re-rendered from previous studies ^19,30^.

#### Lightsheet microscopy

Lightsheet microscopy was performed with a custom lightsheet microscope ^76^ on cubes (5-6 mm) of PFA-fixed *Dasyatis pastinaca* propterygium, optically-cleared using ethyl cinnamate (ECi)^77^, as part of a study on tissue autofluorescence ^78^. The methods are as described there; however, we also explored combinations of different excitation laser lines and detection emission filters to create multi-spectral images (e.g. Fig. 2C). This allowed us to distinguish canals/blood vessels autofluorescence (Ex: 561 nm; Em: 593LP, in red) from other structures, such as NADH autofluorescence from chondrocytes (Ex: 488 nm; Em: 525/50, in cyan).

#### Scanning electron microscopy

Images from backscatter electron microscopy (BSE-SEM: Fig. 1F) and cryo-scanning electron microscopy (cryoSEM: Fig. 2G) of Round stingray cartilage were reanalyzed from previous studies ^18,19,25^.

#### Histology

Histological slides from Round stingray cartilage were reanalyzed from previous works, with methods as described previously ^19,21^. The other species’ tissues examined specifically for this study were prepared as follows:

Devil ray specimens were fixed and stored in 10% neutral buffered formalin. Chondrocranium samples were decalcified in 10% ethylenediaminetetraacetic acid (EDTA) at 37°C, pH 7.4, for 8 days and embedded in paraffin. *Mobula mobular*, *M. thurstoni*, *M. tarapacana* and *M. eregoodoo* samples were cut in 5 µm cross-sections and auto-stained with Hematoxylin and Eosin, as well as Masson’s Trichrome. Slides were then imaged at 20x on a Panoramic Scan slide scanner.

Additional non-decalcified *M. thurstoni* and *M. tarapacana* samples were cut into 50 µm cross-sections, fully dehydrated in methanol, and stained using 0.2 mg/mL fast green in methanol solution after DAPI staining (1:1000). Sections were then cleared in dichloromethane and dibenzylether and imaged using a FLUOVIEW FV4000 Confocal Laser Scanning Microscope.

Copper shark, school shark, blue shark and greeneye spurdog pectoral girdle (coracoid bar) specimens were fixed in 4% PFA and stored in 0.1M PBS. Samples were decalcified in 20% EDTA in PBS at room temperature for 30 days and embedded in paraffin. Serial cross-sections (8μm) were mounted and stained with Masson’s Trichrome (MT). Entire cross-sections were imaged using an Olympus SLIDEVIEW VS200 slide-scanner (Olympus, Version 4.1).

For eagle ray and cownose ray specimens, the jaws were removed and fixed in 4% PFA for 24 hours before being stored in phosphate-buffered saline. Samples were decalcified in 10% EDTA until fully decalcified and embedded in paraffin. Samples were cut into 5 µm cross-sections and auto-stained with Masson’s trichrome. The *A. ocellatus* slide was imaged on a Nikon Ni-E upright microscope. The *R. bonasus* slide was imaged on a Leica Aperio ScanScope CS.

