## Supplementary Figures+Tables for "Cartilage canals in sharks and rays show that blood vessels can exist in mature cartilage without triggering endochondral bone formation"

**Supplementary figures and tables.**

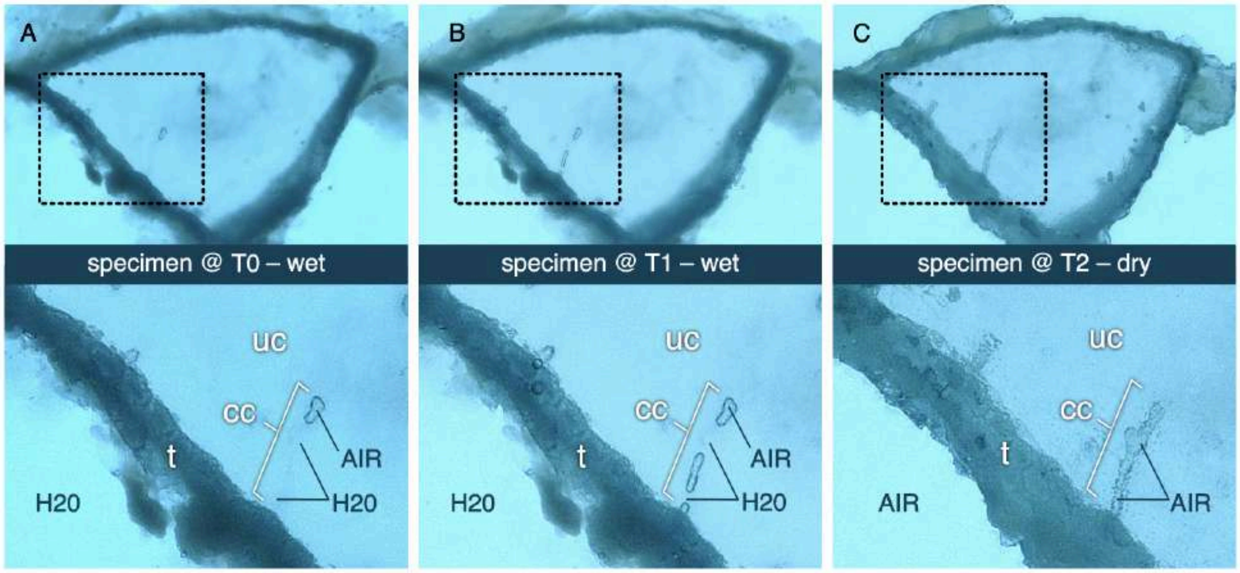


**Figure S1:** Light micrographs showing cartilage canal (cc) visibility changing with dehydration, in a cross section of propterygium from the Round stingray *U. halleri* (11 cm DW). The dashed boxes in the upper row indicate the regions of unmineralized cartilage (uc) and tesserae (t) magnified in the bottom row, shown from left to right over three consecutive time points (T0-2). A) Fresh cut (hydrated) specimen, where a small amount of air has been introduced into the distal end of the canal. B-C) As the specimen dries, air gradually fills the canal back to its proximal end at the tesseral layer, revealing the canal's full extent.

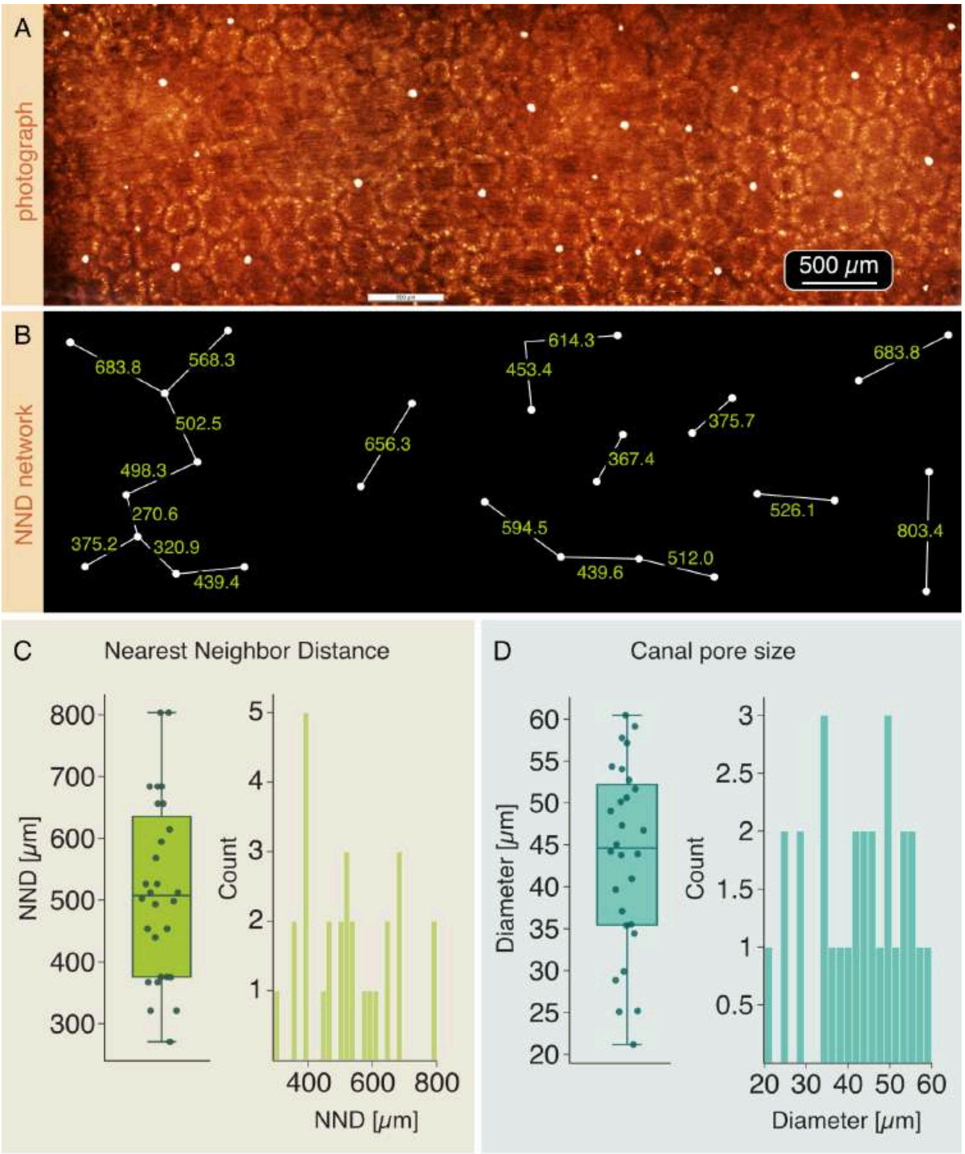


**Figure S2:** Canal pore morphometrics from Round stingray *U. halleri*; A) A backlit light micrograph of tesserae in a flat region of interest of the propterygium, with most of the unmineralized cartilage removed; pores are visible as white spots. B) The isolated network of canal pores showing the calculated nearest neighbor distances (NND, pore edge-to-edge). C-D) Nearest neighbor distances and canal pore size (maximum Feret Diameter) from 28 canal pores. See also Table S1.

**Figure S3:** Elasmobranch canals from those examined species not featured in Figure 4. Arrows indicate cartilage canals. A) *Aetobatus ocellatus*, B) *Hypanus sabinus*, C) *Pteroplatytrygon violacea*, D) *Mobula mobular*, E) *Mobula thurstoni*, F) *Mobula eregoodoo*, G) *Raja binoculata*, H) *Raja asterias*, J) *Raja clavata*, K) *Raja eglanteria*, L) *Rhina ancylostoma*, M) *Narcine bancroftii*, N) *Mustelus canis*, O) *Prionace glauca*, P) *Alopias vulpinus*, Q) *Chiloscyllium plagiosum*. In all images, canals are ~30-50 μm in diameter; panels A,F,L show canals with partial or complete lining by tesserae. The figure includes images of macroscopic skeletal sections (B,C,G,K,M,N,P,Q) and close-up images of individual canals (A,D,E,F,H,J,L,O); canals are sectioned longitudinally in all cases, except G and O. See Table S3 for the particular skeletal elements examined.

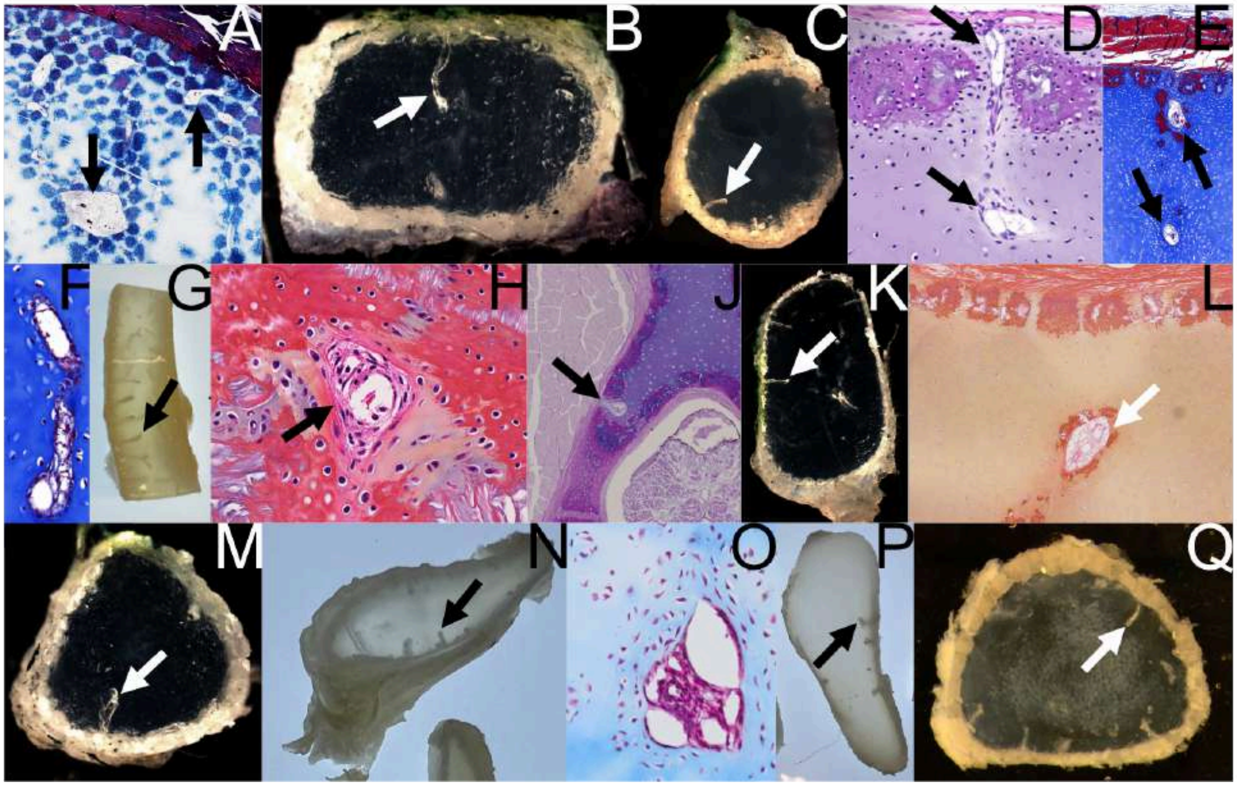

**Table S1:** Canal pore morphometrics, derived from measurements from transmitted light micrographs from a 6.2x2.0mm ROI of *U. halleri* tessellated cartilage (see also Fig. S2).

| | Mean | SD | Min | Max |
|---|---|---|---|---|
| Pore diameter (µm) | 43.6 | 11.1 | 21.2 | 60.5 |
| Nearest neighbor distance (µm) | 515.7 | 144.3 | 270.6 | 803.4 |

**Table S2:** Canal morphometrics, derived from an Amira-based quantification of PTA-stained µCT data of a complete *U. halleri* propterygium (see Fig. 2A,B).

| | Mean | SD | Min | Max |
|---|---|---|---|---|
| Mean radius (µm) | 12.05 | 2.66 | 10.99 | 30.24 |
| Curved length (µm) | 453.29 | 337.89 | 96.14 | 1766.42 |
| Tortuosity | 1.08 | 0.08 | 1 | 1.66 |
| Distance between canals (µm) | 571.42 | 176.79 | 115.13 | 1280.16 |

Total number of canals measured: $n = 408$; Ratio of canal to cartilage volume: 0.00296 (0.296%)

**Table S3:** Elasmobranch samples examined, listed by order and major group (shark or batoid; see Fig. 4). Canal presence is indicated in the fourth column: species are marked as having canals if at least one canal was unequivocally observed across numerous specimens. Subsequent columns list the anatomical location of observations, the study methods used, and literature sources, when data were reanalyzed from previous studies.

| Species | Order | Shark/batoid | Canals? | Location | Methods | Source |
|---|---|---|---|---|---|---|
| *Urobatis halleri* | Myliobatiformes | Batoid | Y | PT[1–5] J[6] | H[1,2], cSEM[6], BSE-SEM[7], P[3], SR-µCT[4], µCT, LS[5] | [1–6] Current study |
| *Aetobatus ocellatus* | Myliobatiformes | Batoid | Y | J | H | BF |
| *Dasyatis pastinaca* | Myliobatiformes | Batoid | Y | PG | H | [5] |
| *Hypanus sabinus* | Myliobatiformes | Batoid | Y | PG[8],J[9,10] | P | [8–10] |
| *Pteroplatytrygon violacea* | Myliobatiformes | Batoid | Y | PG | P | [8,11] |
| *Mobula mobular* | Myliobatiformes | Batoid | Y | CH | H | TH+VCA |
| *Mobula thurstoni* | Myliobatiformes | Batoid | Y | CH | H | TH+VCA |
| *Mobula tarapacana* | Myliobatiformes | Batoid | Y | CH | H | TH+VCA |
| *Mobula eregoodoo* | Myliobatiformes | Batoid | Y | CH | H | TH+VCA |
| *Rhinoptera bonasus* | Myliobatiformes | Batoid | Y | J,PG | H | BF+MND |
| *Leucoraja erinacea* | Rajiformes | Batoid | Y | HY | P | [9,10,12] |
| *Raja binoculata* | Rajiformes | Batoid | Y | HY | P | [9,10] |
| *Raja asterias* | Rajiformes | Batoid | Y | J | H | MDT |
| *Raja clavata* | Rajiformes | Batoid | Y | CH?[13],S[12] | I | [12,13] |
| *Raja eglanteria* | Rajiformes | Batoid | Y | PG | P | [8] |
| *Rhina ancylostoma* | Rhinopristiformes | Batoid | Y | NA | H | [14] |
| *Narcine bancroftii* | Torpediniformes | Batoid | Y | HY | P | [8] |
| *Carcharhinus brachyurus* | Carcharhiniformes | Shark (Galea) | N | PG | H | MYB+SM |
| *Carcharhinus plumbeus* | Carcharhiniformes | Shark (Galea) | Y | HY | P | [9,10] |
| *Galeorhinus galeus* | Carcharhiniformes | Shark (Galea) | Y | PG | H | MYB+SM |
| *Mustelus canis* | Carcharhiniformes | Shark (Galea) | Y | HY | P | [9,10] |
| *Prionace glauca* | Carcharhiniformes | Shark (Galea) | Y | PG | H | MYB+SM |
| *Scyliorhinus canicula* | Carcharhiniformes | Shark (Galea) | N | J,HY,NA,PG | H | [12,15] |
| *Scyliorhinus retifer* | Carcharhiniformes | Shark (Galea) | N | J,HY | P | MND |
| *Alopias vulpinus* | Lamniformes | Shark (Galea) | Y | HY | P | [9,10] |
| *Cetorhinus maximus* | Lamniformes | Shark (Galea) | N? | B | P | [16] |
| *Isurus oxyrinchus* | Lamniformes | Shark (Galea) | Y | HY | P | [9,10] |
| *Chiloscyllium plagiosum* | Orectolobiformes | Shark (Galea) | Y | HY | P | [9,10] |
| *Hexanchus griseus* | Hexanchiformes | Shark (Squalea) | N | HY | P | [9,10] |
| *Notorynchus cepedianus* | Hexanchiformes | Shark (Squalea) | N | HY | P | [9,10] |
| *Somniosus pacificus* | Squaliformes | Shark (Squalea) | N | HY | P | [9,10] |
| *Squalus acanthias* | Squaliformes | Shark (Squalea) | N | HY | P | [9,10] |
| *Squalus chloroculus* | Squaliformes | Shark (Squalea) | N | PG | H | MYB+SM |

| | | | | | | |
|---|---|---|---|---|---|---|
| *Squatina squatina* | Squatiniformes | Shark (Squalea) | Y | HY | P | [9,10] |

<u>*Abbreviation key*</u>*:*

*Anatomical location:* J=Jaws; CH=Chondrocranium; HY=Hyoid arch; NA=Neural arch; PG=Pectoral/pelvic girdle (including propterygium, fin radials); S=synarcual (fused vertebrae)

*Methods:* BSE-SEM=backscatter electron SEM; cSEM=cryo-SEM; H=histology; I=Illustration; LS=Light-sheet microscopy; P=Skeleton section photography; SR-µCT=Synchrotron µCT

*Source:* Literature listed by numbered citation, personal observations listed by author initials: BF=Benjamin Flaum; MDT=Mélanie Debiais-Thibaud; MND=Mason Dean; MYB=Maximus Yeatman-Biggs; SM= Susan McGlashan; TH=Theda Hinrichs; VCA=Victoria Camilieri-Asch

**Table S4:** Non-elasmobranch cartilage canal examples, listed by clade and family (see Fig. 4), with canal presence indicated. Tetrapod vertebrates are far more studied than other groups, with a focus on several particular model species (e.g. chicken, dog, horse, human, mouse). In contrast, literature for non-amniote examples is sparse. A group's designation —especially in the non-tetrapod examples— should therefore not be taken as indication that all members of the group have or do not have canals. Note, in our literature survey, the many published investigations of actinopterygian skeletal histology typically do not mention cartilage canals; the study[17] of the marine teleost *Sparus auratus*, however, specifically notes the lack of canals in the cartilage.

| Clade | Example families | Canals? | Sources |
|---|---|---|---|
| Agnatha | Myxinidae (hagfish) | N | [18–20] |
| Agnatha | Petromyzontidae (lamprey) | N | [21,22] |
| Holocephali | Chimaeridae (chimaera) | N | [12,23] |
| Actinopterygii | Acipenseridae (sturgeon) | Y | [13] |
| Actinopterygii | Sparidae (seabream) | N | [17] |
| Amphibia | Ranidae (frog) | Y | [24] |
| Reptilia | Varanidae (lizard) | Y | [25,26] |
| Reptilia | Dermochelyidae (turtle) | Y | [27] |
| Reptilia | Phasianidae (chicken) | Y | [28–31] |
| Mammalia | Hominidae (human) | Y | [32–35] |
| Mammalia | Muridae (mice) | Y | [36–39] |
| Mammalia | Equidae (horse) | Y | [40–42] |
| Mammalia | Canidae (dog) | Y | [43–45] |
| Mammalia | Leporidae (rabbit) | Y | [39,46–48] |
| Mammalia | Suidae (pig) | Y | [49–51] |
